# CENP-B dynamics at centromeres is regulated by a SUMOylation/ubiquitination and proteasomal-dependent degradation mechanism involving the SUMO-targeted ubiquitin E3 ligase RNF4

**DOI:** 10.1101/245597

**Authors:** Jhony El Maalouf, Pascale Texier, Indri Erliandri, Camille Cohen, Armelle Corpet, Frédéric Catez, Chris Boutell, Patrick Lomonte

**Affiliations:** Univ Lyon, Université Claude Bernard Lyon 1, CNRS UMR 5310, INSERM U 1217, LabEx DEVweCAN, Institut NeuroMyoGène (INMG), team Chromatin Assembly, Nuclear Domains, Virus. F-69100, Lyon, France; Univ Lyon, Université Claude Bernard Lyon 1, CNRS UMR 5286, INSERM U 1052, Centre de Recherche en Cancérologie de Lyon. F-69000, Lyon, France; MRC-University of Glasgow Centre for Virus Research, Glasgow G61 1QH, Scotland (UK)

## Abstract

Centromeric protein B (CENP-B) is a major constituent of the centromere. It is a DNA binding protein that recognizes a specific 17-nt sequence present in the centromeric alphoid satellite repeats. CENP-B importance for centromere stability has only been revealed recently. In addition to its DNA binding properties, CENP-B interacts with the histone H3 variant CENP-A and CENP-C. These interactions confer a mechanical strength to the kinetochore that enables accurate sister chromatids segregation to avoid aneuploidy. Therefore, understanding the mechanisms that regulate CENP-B stability at the centromere is a major unresolved issue for the comprehension of centromere function. In this study, we demonstrate that lysine K402 of CENP-B is a substrate for SUMO post-translational modifications. We show that K402 regulates CENP-B stability at centromeres through a SUMOylation/ubiquitination and proteasomal-dependent degradation mechanism involving the SUMO-Targeted Ubiquitin E3 Ligase RNF4/SNURF. Our study describes SUMOylation of CENP-B as a major post-translational modification involved in centromere dynamics.

## Introduction

Since its discovery as one of the three major centromere-associated antigens recognized by autoimmune sera of patients with scleroderma (Earnshaw and Rothfield, 1985; Earnshaw et al., 1987a), centromeric protein B (CENP-B) has turned out to be a very complex and multifunctional protein. CENP-B is conserved among species from yeast to human, highlighting its functional importance, although not essential for mouse development until birth (Hudson et al., 1998; Kapoor et al., 1998; Perez-Castro et al., 1998). CENP-B exists as a homodimer that binds to DNA through a specific 17-bp motif (termed the CENP-B box) present in the alphoid repeated sequences constituting the DNA backbone of the centromere region of all human and mouse chromosomes, with the exception of the Y chromosome (Masumoto et al., 1989; Muro et al., 1992; Yoda et al., 1992; Kitagawa et al., 1995; Tawaramoto et al., 2003). Although not required for the maintenance of the mouse and Y chromosomes during cell divisions, CENP-B and the CENP-B box were shown to be essential for the *de novo* assembly of a functional centromere using artificial human chromosomes (Masumoto et al., 1998; Ohzeki et al., 2002). Accordingly, CENP-B knockout or depleted cells show a higher rate of chromosome mis-segregation and chromosomal instability (Fachinetti et al., 2015). In addition, CENP-B null mice show reproductive and developmental dysfunctions due to defects in the morphogenesis of the high *Cenpb*-gene expressing and highly mitotically active uterine epithelial tissues (Fowler et al., 2000; Fowler et al., 2004). At the molecular level, CENP-B accumulation at centromeres has been linked to its capacity to interact with the N-terminal tail of the centromere-specific histone H3 variant CENP-A (Fachinetti et al., 2013). The sustained upkeep of CENP-B at the centromere through its interaction with CENP-A contributes to the stability of chromosome segregation, conferring a major role to the CENP-B/CENP-A tandem as stabilizers of the kinetochore nucleation (Fachinetti et al., 2013). In addition, CENP-B also directly interacts with the C-terminal tail of CENP-C, an essential protein for kinetochore assembly that independently interacts with CENP-A, which contributes to the functional stabilization of the kinetochore (Fachinetti et al., 2015; Hoffmann et al., 2016).

Apart of an increased residence time at the centromere during the G1/S phase of the cell cycle, little is known about the dynamic regulation of CENP-B at the centromere (Hemmerich et al., 2008). In the present study we investigated the role of SUMO (Small Ubiquitin Modifier) modification on CENP-B stability at centromeres. We demonstrate that CENP-B could be SUMOylated in vitro on several lysine (K) residues, with K402 being the major SUMO modified residue. CENP-B K402 modification influenced CENP-B turnover at centromeres through a proteasome-dependent degradation mechanism involving the SUMO-Targeted Ubiquitin Ligase (STUbL) RNF4/SNURF (hereafter called RNF4).

## Results

### CENP-B is SUMOylated in vitro and in vivo

Recently published SUMO’omics studies detected CENP-B as a potential substrate for SUMOs modifications (Bruderer et al., 2011; Hendriks et al., 2015; Sloan et al., 2015). This implies that CENP-B post-translational modifications by SUMO can have a major role in the regulation of CENP-B dynamics at centromeres, and consequently on the centromeres activity itself. To detect lysine residues present in putative SUMOylation consensus sites we analyzed *in silico* the human CENP-B sequence using the JASSA (Joint Analyzer of SUMOylation Site and SIMs) software (Beauclair et al., 2015). Four potential SUMOylation sites were found, three of which (K28, K58, K76) associated with the N-terminal DNA binding (DBD) domain and one (K402) present in the core of the protein (Fig. 1A). To determine if any of the 4 K residues were acceptor sites for CENP-B SUMOylation, point mutations were introduced in the CENP-B nucleotide sequence to produce CENP-B proteins mutated on each K individually or in combination (Fig. 1B). We did not use the usual modification K>R but rather K>G (or P) because due to the high GC content of the CENP-B nucleotide sequence (over 65%) some SUMOylation site-associated Ks where not in a proper environment to design oligonucleotides suitable to perform K>R site mutagenesis. *In vitro* SUMOylation assays showed that CENP-Bwt and CENP-B_3K were SUMOylated to comparable levels to that of the positive control promyelocytic leukemia (PML) protein (Fig. 1C, compare lanes 4, 6, and 10). In contrast, CENP-B_4K had a significantly reduced SUMOylation pattern (Fig. 1C, lane 8). This result confirms that CENP-B can be SUMOylated and demonstrates that K402 is the main SUMOylated lysine. Quantitation of unmodified forms of CENP-Bwt and CENP-B_3K using the LI-COR Infrared Fluorescent Imaging System suggested that other K in the DBD could be SUMO modified (Fig. S1A). We performed additional assays using a CENP-B protein deleted of the first 129 aa constituting the DBD domain (CENP-BΔDBD; (Yoda et al., 1992; Tanaka et al., 2005b) or the same protein with the additional mutation of K402 (CENP-BΔDBD_K402G) (Fig. 1D and S1B). Compared to CENP-Bwt, CENP-BΔDBD SUMO signal was substantially decreased (compare lanes 2 and 6), and CENP-BΔDBD_K402G lost almost the entire SUMOylation signal (compare lanes 2 and 4). This confirms that although K402 contributes for the majority of the CENP-B SUMO signal, several lysine residues in the DBD are probably also substrates for SUMO modifications. To demonstrate that CENP-B could be covalently linked to SUMO *in vivo*, we performed denaturing HIS-pull down assays from cells ectopically expressing CENP-Bwt and 6xHIS-tagged SUMO-1. Although the HA tag has, on its own and unspecifically, an affinity for the Ni-NTA agarose resin, the result clearly showed that a high molecular weight extra CENP-B band was reproducibly detected when CENP-B was co-expressed with 6xHIS-SUMO-1 (Fig. 1E). This result indicates that CENP-B could be covalently linked to SUMO in cells, although this is likely to represent a minute part of the CENP-B protein. Altogether, these data demonstrate that CENP-B is a substrate for SUMOylation, which validates the previously published SUMO’omics data.

**Figure 1.**
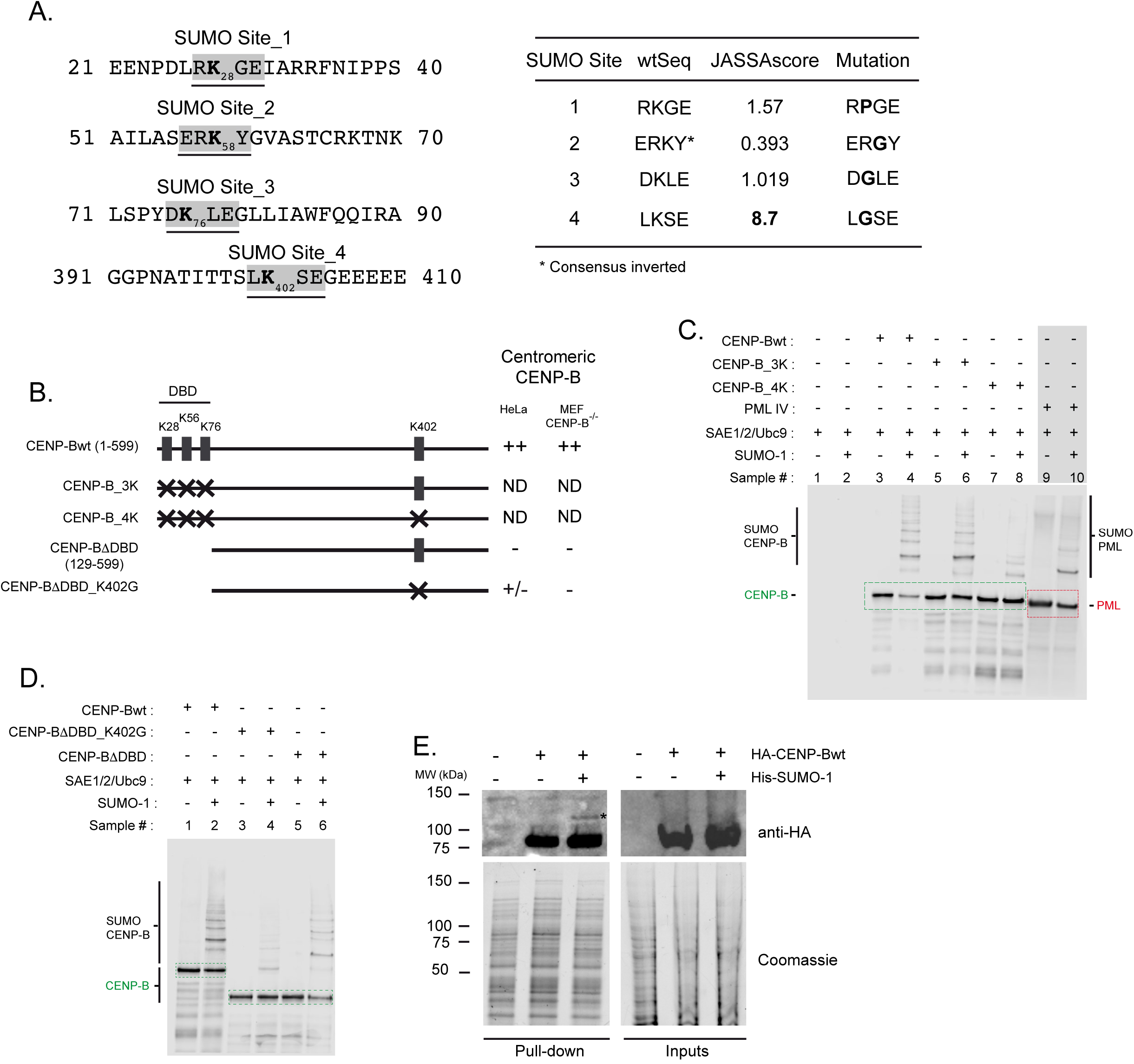
Centromeric protein B (CENP-B) is SUMOylated in vitro and in vivo (A) Left: Consensus sequences for SUMOylation sites in the CENP-B sequence. Right: Joined Advanced Sumoylation Site and SIM Analyser (JASSA; http://www.jassa.sitesgb.info/) (Beauclair et al., 2015) scores for potential SUMOylated K and mutations performed for each K (for details see the Materials and Methods section). * indicates a consensus inverted site. (B) Left: Schematic representations of the different tagged CENP-B proteins used in this study. Right: Summary of the immunofluorescence data shown in Figs. 2 and Table 1 for the colocalization of ectopically expressed CENP-B proteins with centromeres in HeLa and MEF CENP-B−/− cells. ++: all CENP-B spots co-localize with centromeres; +/−: a subset of CENP-B spots co-localizes with centromeres; −: no colocalization of CENP-B with centromeres. (C) In vitro SUMOylation assay of 6xHis-FLAG-CENP-B, 6xHis-FLAG-CENP-B_3K, and 6xHis-FLAG-CENP-B_4K. 6xHis-FLAG-PML isoform IV was used as a positive control. WB was performed using an anti-FLAG antibody to detect tagged proteins. The LI-COR Infrared Fluorescent Imaging System was used to detect and quantify the signals (see original blot in Fig. S1A). Boxes: unmodified CENP-B or PML. (D) In vitro SUMOylation assay of 6xHis-FLAG-CENP-B, 6xHis-FLAG-CENP-BΔDBD, and 6xHis-FLAG-CENP-BΔDBD_K402G. Up: Western Blot (WB) was performed using an anti-FLAG antibody to detect tagged proteins. LI-COR was used to detect and quantify signals (see original blot in Fig. S1B). Boxes: unmodified CENP-B. (E) His pull-down assays using extracts from cells co-transfected with expression vectors for HA-CENP-B and 6xhis-SUMO-1. WB was performed using an anti-HA antibody to detect CENP-B proteins pulled-down with His-SUMO-1. * indicates an extra CENP-B band associated with the co-expression of CENP-B with His-SUMO-1.

**Figure 2.**
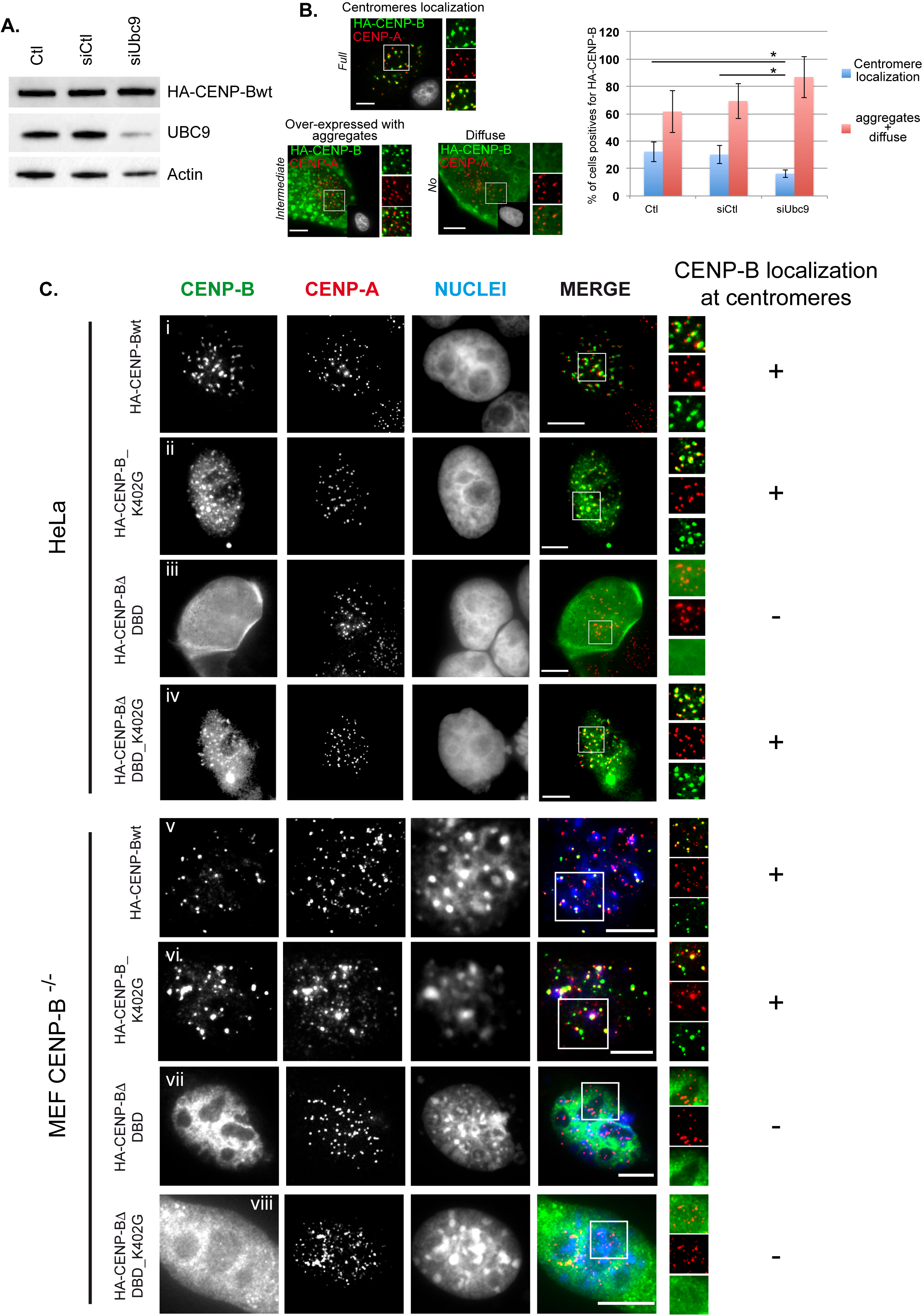
CENP-B targeting at centromeres is dependent on SUMOylation but not CENP-B K402 (A) WB of HeLa cell extracts not transfected (Ctl) or transfected with control siRNA (siCtl) or Ubc9 siRNA (siUbC9) for 48 h prior to transfection with HA-CENP-B expressing plasmid. Decreases in Ubc9 induced by siUbc9 do not affect the overall expression of HA-CENP-B. Actin was used as a loading control. (B) Left: Immunofluorescence to detect HA-CENP-B patterns in HeLa cells not transfected (Ctl) or transfected with control siRNA (siCtl) or Ubc9 siRNA (siUbC9). Three major patterns were observed for the HA-CENP-B signal: (i) centromeres localization; (ii) over-expressed with aggregates; (iii) diffuse. Each pattern shows different frequencies of co-localization between HA-CENP-B and CENP-A (full, intermediate, or no-colocalization). Right: Quantification of the patterns in cells positives for HA-CENP-B signal and in the different samples. * indicates significant difference (p value ≤ 0.05) in the scores based on Student’s *t*-test. Bars = 5 μm. (C) Immunofluorescence performed on HeLa or MEF CENP-B−/− cells ectopically expressing wild type CENP-B or several CENP-B mutants. +: indicates that co-localization with CENP-A could be observed; −: indicates that no co-localization with CENP-A could be observed. Bars = 5μm.

### SUMO pathway but not K402 is essential for CENP-B targeting to the centromere

A previous study showed the exchange of CENP-B at centromeres to be highly dynamic during the G1-S phases of the cell cycle (Hemmerich et al., 2008). To determine if SUMOylation was involved in CENP-B targeting to centromeres, we analyzed the accumulation of ectopically expressed HACENP-Bwt at centromeres in cells depleted of Ubc9, the only known mammalian E2 SUMO conjugating enzyme, by RNA interference (Fig. 2A and B). Depletion of Ubc9 did not affect HA-CENP-B expression (Fig. 2A), but significantly influenced its targeting to centromeres in cells showing a positive signal for HA-CENP-B (32.3 ± 7.2%, 30.2 ± 6.5% vs 16.3 ± 2.8% in Ctl (nosiRNA), siCtl and siUbc9 treated cells, respectively) (Fig. 2B). These data suggest that Ubc9 activity, and hence a functional SUMOylation pathway, is required to target CENP-B to centromeres. To determine if K402 was involved in the targeting of CENP-B to centromeres, CENP-Bwt or CENP-B_K402G were ectopically expressed in HeLa cells. Co-localization with CENP-A of exogenous CENP-B wt and CENP-B_K402G was analyzed by immunocytochemistry. No major changes in CENP-B targeting at centromere was detected (Fig. 2Ci and ii). Only the CENP-BΔDBD showed a complete absence of CENP-B at centromeres (Fig. 2Ciii), as expected from previous studies (Pluta et al., 1992) and seemingly easily explained by its inability to bind to CENP-B boxes. However, and strikingly, the additional mutation of K402 in the CENP-BΔDBD background restored to some extent a CENP-B centromeric signal (Fig. 2Civ and Fig. 3Bv). Endogenous CENP-B is present in HeLa cells, potentially interacting with the ectopic CENP-B through the C-terminal dimerization domain and introducing a bias of targeting. To avoid any misinterpretation due to dimerization with endogenous CENP-B, we performed similar experiments in mouse embryonic fibroblasts (MEF) isolated from CENP-B knock out mice in which both CENP-B alleles have been inactivated (MEF CENP-B^−/−^) (Kapoor et al., 1998; Okada et al., 2007). Although the ectopic CENP-B is of human origin its non-mutated form was accurately targeted to mouse centromeres (Fig. 2Cv and Table 1). Mutation of K402 alone did not modify the targeting of CENP-B to centromeres compared to the CENP-Bwt. Consistently, a slight increase of the CENP-B_K402G signal at centromere was observed (Fig. 2Cvi and Table 1). Like in HeLa cells the CENP-BΔDBD was completely diffuse in the nucleoplasm with no co-localization with centromeres (Fig. 2Cvii and Table 1). Unlike in HeLa cells the CENP-BΔDBD_K402G showed the same pattern as CENP-BΔDBD with no accumulation of CENP-BΔDBD_K402G signal at centromeres (Fig. 2Cviii and Table 1). These data suggest that although K402 is not involved in the targeting of CENP-B to centromeres, it could potentially impact on the CENP-B residence time.

**Figure 3.**
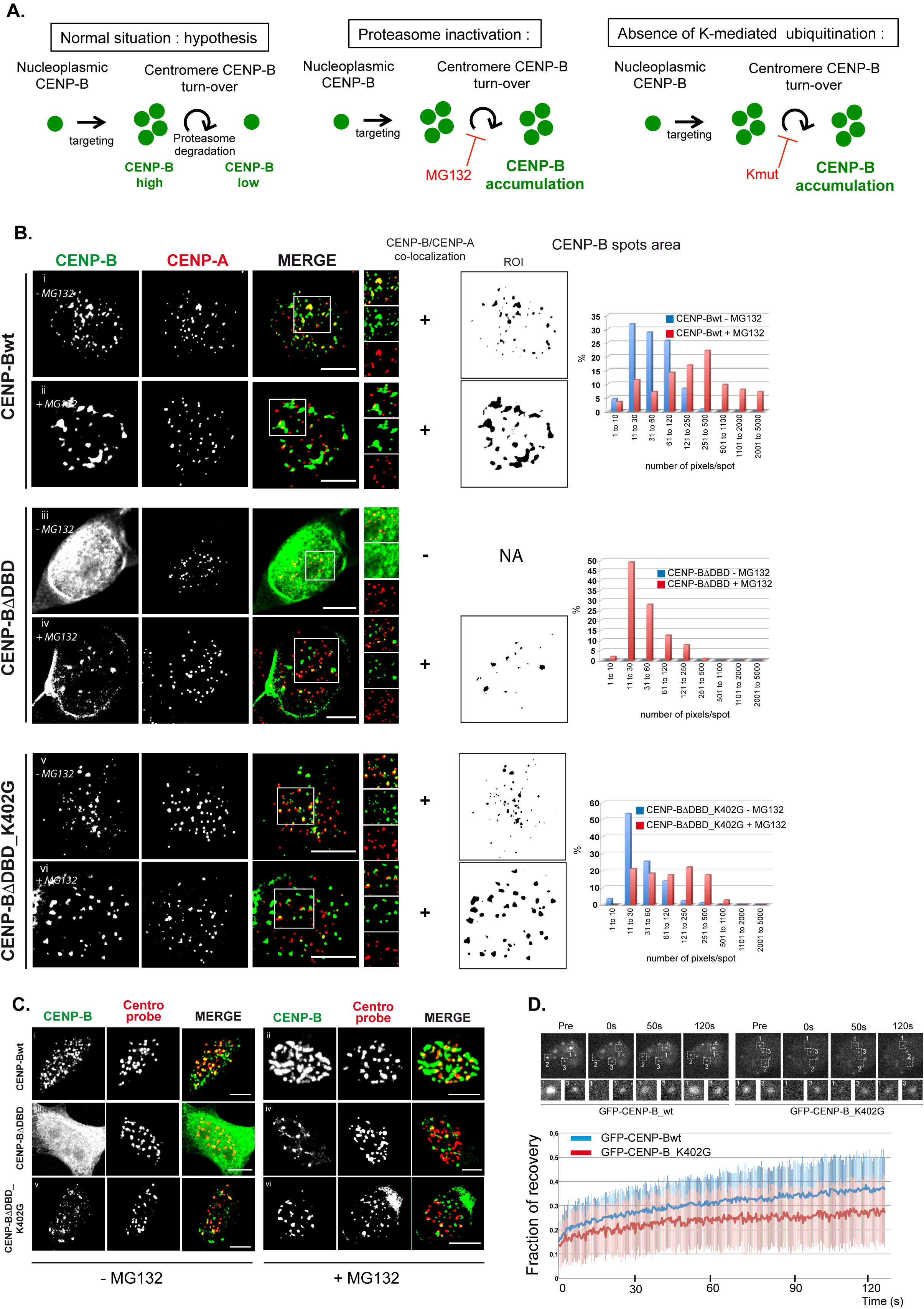
CENP-B accumulation at centromeres and CENP-B turnover are affected by proteasome inactivation and mutation of K402. (A) Synopsis of the potential regulation of CENP-B amount at centromeres. (B) Left: Immunofluorescence to detect ectopically expressed HA-CENP-Bwt, HA-CENP-BΔDBD, and HA-CENP-BΔDBD_K402G in HeLa cells in the absence or presence of the proteasome inhibitor MG132. Bars = 5μm. +: indicates co-localization between CENP-B and CENP-A was observed; −: indicates that no co-localization between CENP-B and CENP-A was observed. Middle: Size of the CENP-B spots according to the region of interest (ROI) as determined using ImageJ software. HA-CENP-BΔDBD does not show focused signals (spots) above the general nucleoplasmic background. Right: Quantification of the size of the spots (pixel/spots) in different experimental conditions. Data are means of three independently performed experiments. NA= Not applicable. (C) Immuno-FISH for the detection of ectopically expressed HA-CENP-Bwt, HA-CENP-BΔDBD, and HA-CENP-BΔDBD_K402G in HeLa cells in the absence or presence of the proteasome inhibitor MG132. Centromeres were detected using a human chromosome pan-centromeric biotin probe. Bars = 5 μm. (D) Fluorescence Recovery After Photobleaching (FRAP) analysis of GFP-CENP-Bwt and GFP-CENP-B_K402G mobility at centromeres. Top: representative images showing the centromere-associated fluorescence before, during (0 s) and after (50 and 120 s) photobleaching. Boxed areas 1 and 2 represent two centromeres simultaneously bleached; area 3 represents an unbleached centromere. Bottom: measure of the fluorescence recovery for GFP-CENP-Bwt and GFP-CENP-B_K402G.

### Proteasome inhibition and mutation of K402 induces CENP-B accumulation at centromeres

The major contribution of CENP-B K402 in CENP-B SUMOylation *in vitro* suggests a specific implication of K402 in CENP-B dynamics. Lysine SUMOylation has been shown to influence protein stability through SUMO targeted ubiquitination and proteasomal-degradation (Tatham et al., 2008). To determine if CENP-B SUMOylation influenced its stability at centromeres the accumulation of ectopically expressed CENP-Bwt, CENP-BΔDBD, and CENP-BΔDBD_K402G were measured in HeLa cells in the presence or not of the proteasome inhibitor MG132 (Fig. 3A). In the presence of MG132, CENP-Bwt showed a clear aggregation pattern with most of the CENP-B signal co-localized with or juxtaposed to centromeres. Some CENP-B aggregates were also visible not-colocalizing with centromeres possibly highlighting neo-synthesized CENP-B not incorporated in centromeres and not degraded (Fig. 3Bi and ii left). Overall, the addition of MG132 led to an increase in the average size of CENP-B spots (Fig. 3Bi and ii middle (ROI)) represented by a number of pixels/spot that shifted towards higher values (Fig. 3B up-right). CENP-BΔDBD and CENP-BΔDBD_K402G lack the DBD domain and are not able to bind the CENP-B boxes of the alphoid sequences. Accordingly, no CENPA co-localization was observed with the diffuse CENP-BΔDBD nucleoplasmic signal in the absence of MG132 (Fig. 3Biii left, and middle). The addition of MG132 partially restored CENP-BΔDBD colocalization with CENP-A (Fig. 3Biv left, and middle (ROI)) and enabled the detection of spots with low values of pixels/spot (Fig. 3B middle-right). Strikingly, the sole mutation of K402 within the CENP-BΔDBD background was enough to partially restore CENP-B signal at centromeres, a phenotype that was amplified in the presence of MG132 (Fig. 3Bv and vi left, and middle (ROI)). Similarly to CENP-Bwt, the number of pixels/spot shifted towards higher values in the presence of MG132 (Fig. 3B down-right). Ectopically expressed CENP-Bwt showed very large aggregates in the presence of MG132 most of them co-localized/juxtaposed to CENP-A signal. The enlargement of the CENP-B signal could be due to its oligomerization and/or to a modification of the shape of the CENP-B box containing, but CENP-A deficient, centromere regions not necessarily detectable by the sole labeling of CENP-A. To verify the latter, immuno-FISH was performed to detect CENP-B and centromeres using a probe labelling as many centromere sequences as possible (Fig. 3C). Data showed that the size of the signal labelled by the probe was not enlarged in cells expressing ectopic CENP-Bwt and treated with MG132. This result does not favor an unwinding of the centromere regions due to the presence of undegraded CENP-B. Overall, the results show that proteasome inhibition induces the aggregation of CENP-B at or in the immediate vicinity of centromeres, which highly suggests a control of CENP-B stability by the proteasome with the contribution of K402.

CENP-BΔDBD, and CENP-BΔDBD_K402G lose their capacity to bind to the CENP-B boxes present in the alphoid sequences. However, the C-terminal region of CENP-B is involved in its dimerization at the centromeres. Therefore, CENP-BΔDBD, and CENP-BΔDBD_K402G could be retained at the centromeres in a stochastic manner through the interaction with endogenous CENP-B in HeLa cells. If the process of degradation of the mutated CENP-B is faster than its residence time, the centromeric accumulation of the protein by the sole interaction through its C-terminal domain is likely to be too weak to be visualized. In agreement with this hypothesis, CENP-BΔDBD was only visualized localizing with centromeres in cells treated with the proteasome inhibitor MG132 (compare Fig. 3Biii and iv). The detection of CENP-BΔDBD_K402G at centromeres, even in the absence of MG132 treatment, suggests that K402 could be involved in the turnover of CENP-B at centromeres in a proteasome-dependent manner. If so, the exchange rate of CENP-BΔDBD_K402G at centromeres would be expected to decrease in comparison to wild-type CENP-B. To test this hypothesis, we performed fluorescent recovery after photobleaching (FRAP) experiments using GFP-CENP-Bwt and GFP-CENP-B_K402G (Fig. 3D). Both GFP-fused proteins were correctly targeted to centromeres (Fig. S2Ai and ii). Spots of GFP-CENP-B were bleached and recovery of the signals was measured at time points where CENP-B showed its highest rate of mobility (Hemmerich et al., 2008). Data were obtained from at least 4 independent experiments representing over 50 centromeres. The mutation of K402 reduced the recovery of CENP-B at centromeres, indicating that mutated CENP-B had a longer residency time at centromeres. This implied a role of K402 in the exchange rate of CENP-B. Accordingly, CENP-B proteins unable to bind to the CENP-B box, and hence poorly targeted to centromeres, would be expected to have similar dynamics. Accordingly, FRAP analysis of GFP-CENP-BΔDBD and GFP-CENP-BΔDBD_K402G demonstrated that these two mutants had similar levels of mobility (Fig. S2Aiii and iv, and S2B). These data indicate that K402 is indeed implicated in the dynamics of CENP-B localized at centromeres.

### Proteasome inhibition or K402 mutation induce SUMO and ubiquitin accumulation at centromeres

The above data demonstrate that both proteasome activity and K402 mutation influence CENP-B turnover at centromeres. This could simply suggest that CENP-B stability depends on K402 ubiquitination. However, our *in vitro* data showed that K402 takes a major part in the SUMOylation of CENP-B (Fig. 1C and D), suggesting a combined role for SUMOylation and ubiquitination in the regulation of CENP-B stability at centromeres (Fig. 4A). If true, then SUMO and ubiquitin signals should accumulate at centromeres in the presence of CENP-B following proteasome inhibition. Treatment of cells with MG132 showed an increase in SUMO-1 and SUMO-2/3 signals at centromeres in a subset of cells (approximately 10% of interphase cells) (Fig. 4Bi to vi). Ectopically expressed CENP-Bwt, CENP-BΔDBD, and CENP-BΔDBD_K402G co-localized with SUMO-1 at centromeres in MG132-treated (Fig. 4Cii, iv, vi), but not untreated (Fig. 4Ci, iii, v) cells. Similar data were obtained with SUMO-2/3 (Fig. S3). Co-localization of ubiquitin together with CENP-Bwt, CENP-BΔDBD, and CENP-BΔDBD_K402G was also observed at centromeres in MG132-treated cells (Fig. 4D). The remaining accumulation of SUMOs and ubiquitin at centromeres with the CENP-BΔDBD_K402G that lacks the four lysines initially determined as present in a strong SUMO consensus site is not unexpected for at least three reasons: (i) other K present in the CENP-B sequence could be the target of SUMOylation (Bruderer et al., 2011; Hendriks et al., 2015; Sloan et al., 2015); (ii) CENP-Bwt and its mutants could interact with protein that are themselves SUMOylated and/or ubiquitinated; and (iii) because of the presence of endogenous CENP-B and its capacity to dimerize with the ectopic CENP-B it could not be excluded that a subset of endogenous SUMOylated and/or ubiquitinated CENP-B accumulates at centromeres in those conditions. To confirm that SUMO and ubiquitin accumulation at centromeres was directly linked to the accumulation of ectopic CENP-B at centromeres following proteasome inhibition, and not a consequence of the treatment or of the presence of endogenous CENP-B, MEF CENP-B^−/−^ cells were transfected to express CENP-Bwt, CENP-BΔDBD, or CENP-BΔDBD_K402G under MG132 treatment or not. CENP-Bwt was readily observed to accumulate at centromeres in treated and untreated cells (Fig. S4Ai and ii). Like in HeLa cells, colocalization with SUMO-1 (Fig. 4Ei and ii), SUMO-2/3 (Fig. S4Bi and ii) and ubiquitin (Fig. S4Ci and ii) was only observed following MG132 treatment. In contrast, CENP-BΔDBD and CENP-BΔDBD_K402G, which, unlike in HeLa cells, fail to accumulate at centromeres irrespective of the MG132 treatment (Fig. S4Aiii to vi), did not induce the characteristic spotty pattern of centromeric accumulation of SUMO-1 (Fig. 4Eiii to vi), SUMO-2/3 (Fig. S4Biii to vi), and ubiquitin (Fig. S4Ciii to vi). These data demonstrate that SUMO and ubiquitin conjugates stably accumulate at centromeres concomitantly with the recruitment of CENP-B and only if the proteasome is inactive, suggesting that both SUMOylation and ubiquitination contribute to the dynamics of CENP-B at centromeres.

**Figure 4.**
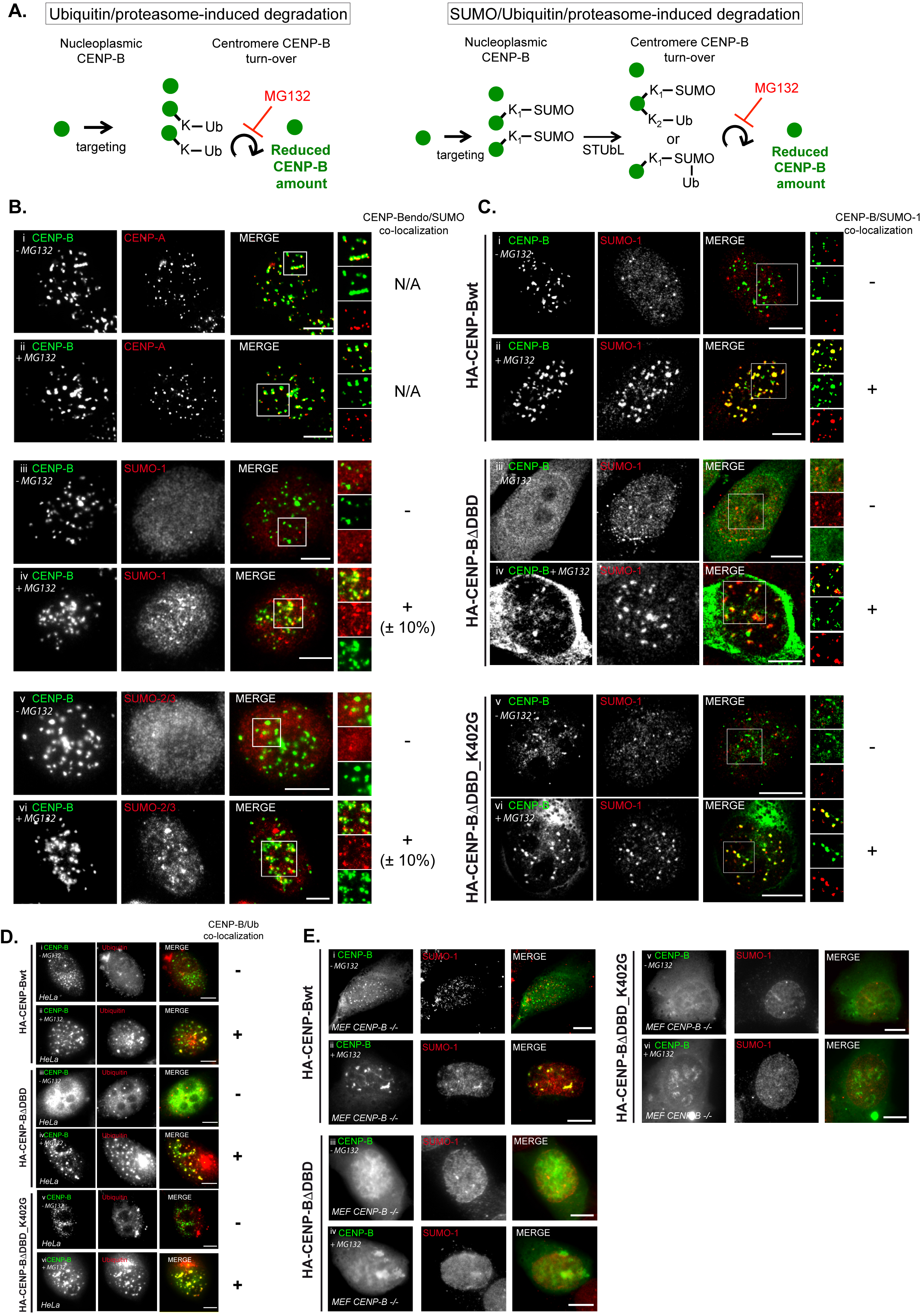
Proteasome inhibition induces SUMO and ubiquitin accumulation at centromeres in a manner dependent on CENP-B targeting at the centromere. (A) Synopsis of the potential regulation of CENP-B amount at centromeres through ubiquitination or SUMOylation/ubiquitination and proteasomal degradation pathways. (B) Immunofluorescence to detect endogenous SUMO-1 and SUMO-2/3 at centromeres (detected by endogenous CENP-B and/or CENP-A), in HeLa cells not treated or treated with the proteasome inhibitor MG132 for 5 h. Bars = 5 μm. N/A: not applicable; +: indicates that colocalizations could be observed; −: indicates that no co-localization could be observed. (C) Immunofluorescence to detect SUMO-1 at centromeres in HeLa cells expressing HA-CENP-Bwt, HA-CENP-BΔDBD, or HA-CENP-BΔDBD_K402G in the absence or presence of the proteasome inhibitor MG132. Bars = 5 μm. +: indicates that co-localization was observed; −: indicates that no co-localization was observed. (D) Immunofluorescence to detect ubiquitin at centromeres in HeLa cells expressing HA-CENP-Bwt, HA-CENP-BΔDBD, or HA-CENP-BΔDBD_K402G in the absence or presence of the proteasome inhibitor MG132. Bars = 5 μm. +: indicates that co-localization was observed; −: indicates that no co-localization was observed. (E) Immunofluorescence to co-detect CENP-B and SUMO-1 at centromeres in MEF CENP-B^−/−^ cells expressing HA-CENP-Bwt, HA-CENP-BΔDBD, or HA-CENP-BΔDBD_K402G in the absence or presence of the proteasome inhibitor MG132. Bars = 5 μm.

### RNF4 STUbL activity regulates CENP-B accumulation at centromeres

RNF4 is an E3 ubiquitin ligase with SUMO-targeting properties that regulates the stability of many SUMO-modified substrates (Tatham et al., 2008). As the stability of CENP-B at centromeres was influenced in both SUMO-and ubiquitination-dependent manner, we investigated the potential role of RNF4 in the turnover of CENP-B at centromeres. Over-expression of RNF4 induced the depletion of CENP-B from centromeres in a subset of cells (48.7 ± 3.2 %, 400 cells counted over five experiments) (Fig. 5Ai and ii; Fig. 5B), but not CENP-A (Fig 5Av), in a manner that was dependent on the catalytic activity of the RING-finger domain of RNF4 (RNF4_DN_) (3.5 ± 3.3 %, p = 1.6 10^−6^) (Fig. 5Aiii and iv; Fig 5B). Interestingly, mutation of the RNF4 RING-domain also increased the number of spots of RNF4 co-localizing with CENP-B or CENP-A at centromeres (Fig 5Avi and vii). Co-transfection of SUMO-1 or SUMO-2 with RNF4wt induced a small but statistically significant increase in cells showing CENP-B depletion (55.8 ± 5.6 %; p = 0.043 for SUMO-1, and 54.2 ± 5.5 %, p = 0.05 for SUMO-2; 500 cells counted over 4 experiments) (Fig. 5B), which suggested a role of SUMOylation in addition to RNF4 in the control of CENP-B stability at centromeres. Co-immunoprecipitation assays using ectopically expressed proteins highlighted an interaction between CENP-B and RNF4 (Fig. 5C). The interaction was likely dependent on SUMOylation because the addition of the SUMO proteases inhibitor N-ethylmaleimide (NEM) was required to maintain this interaction whenever RNF4 (Fig. 5D, two upper gels compare lanes 4 and 5, and 6 and 7) or CENP-B (Fig 5E, upper gel, compare lanes 4 and 5, and 6 and 7) was immunoprecipitated. Addition of NEM also enabled the detection of high molecular weight (HMW) signals detected by the FLAG antibody (Fig. 5D, long exposure membrane). Stripping and reprobing the membranes enabled the detection of SUMOs (Fig. 5D and 5E, two middle gels) and ubiquitin (Fig. 5D, lower gel) moieties. This indicated the likelihood of SUMOs and ubiquitin-modified CENP-B co-IPed with RNF4, although we cannot rule out the possibility of a contribution of other SUMOylated and ubiquitinated proteins than CENP-B and associated with RNF4 to the high MW signals observed. Co-IPs performed in RNF4_DN_ expressing cells showed a reduced ubiquitin signal although SUMO signals remained unchanged, which confirmed the reduced ubiquitination activity of RNF4_DN_, accounting for the lack of effect of RNF4_DN_ on CENP-B stability at centromeres. The amount of putatively SUMO and ubiquitin modified CENP-B co-IPed with RNF4 represented a minute part of the total amount of CENP-B, which may reflect the low amount of CENP-B harboring such modifications in those experimental conditions. WBs performed in cells co-expressing ectopic CENP-B and either RNF4 or RNF4_DN_ showed that RNF4 induced a slight but reproducible decrease of CENP-B that was not observed with RNF4_DN_, suggesting an RNF4-induced and ubiquitin/proteasome-dependent CENP-B degradation (Fig. 5F, up and middle for quantifications). RT-qPCR analyses ruled out any misinterpretation due to a decrease of the FLAG-CENP-B mRNA (Fig. 5F, down).

**Figure 5.**
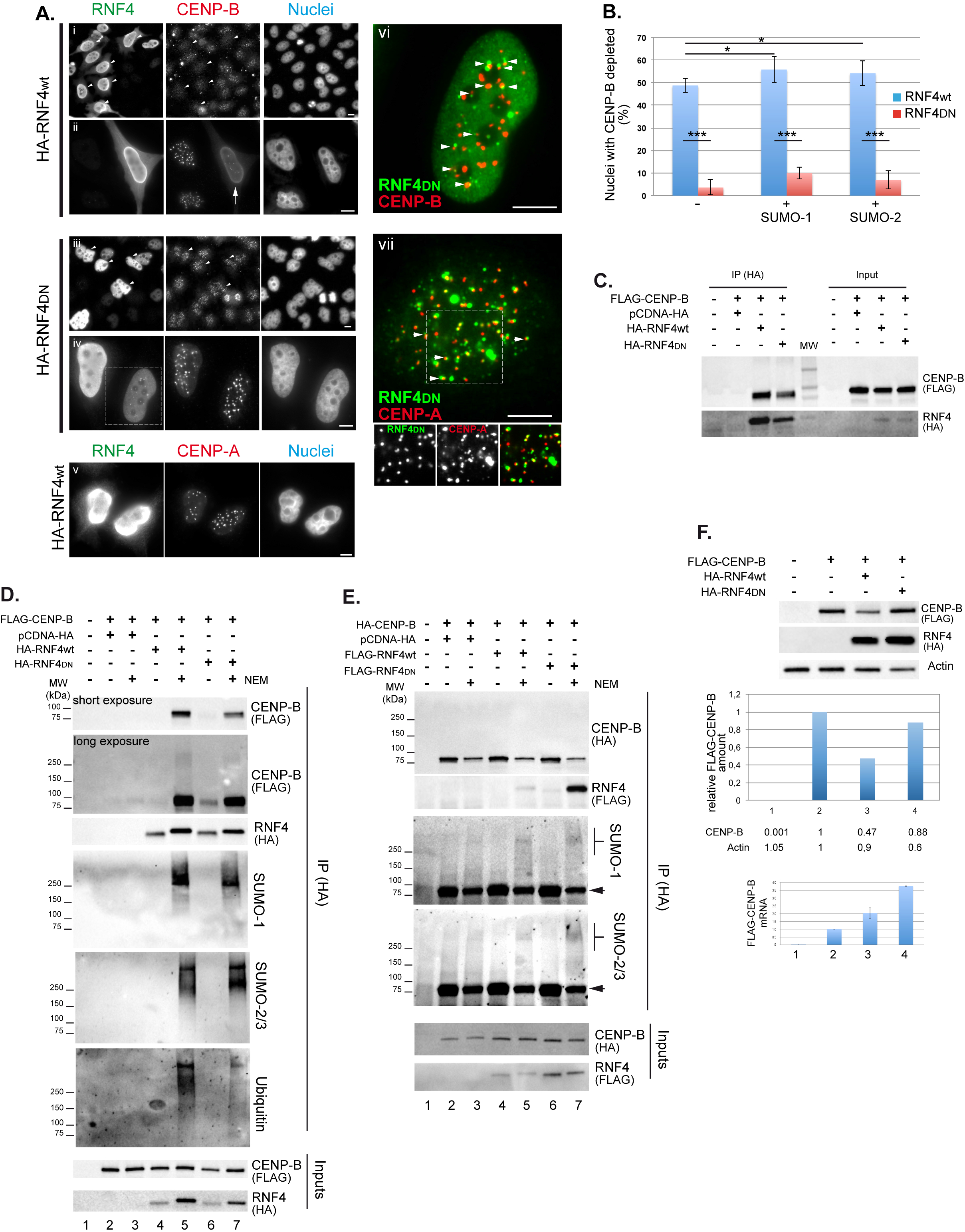
CENP-B stability depends on RNF4 E3 ubiquitin ligase activity (A) Immunofluorescence to detect endogenous CENP-B (i to iv and vi) or CENP-A (v and vii) in HeLa cells ectopically expressing HA-RNF4wt (i, ii, and v) or its dominant-negative (DN) mutant HA-RNF4_DN_ (iii, iv, vi and vii). Two magnifications are shown for CENP-B to better illustrate the effects. Arrows indicate cells expressing HA-RNF4wt or DN and showing a decrease in CENP-B signal or not, respectively. vi and vii illustrate the co-localization of RNF4_DN_ with CENP-B and CENP-A, respectively. Bars = 5 μm. (B) Quantification of cells showing a decrease of CENP-B at centromeres in HeLa cells ectopically expressing HA-RNF4wt or HA-RNF4_DN_ alone or in combination with SUMO-1 or SUMO-2. The results show means of three independent experiments (± standard deviation; SD). * p ≤ 0.05, *** p ≤ 0.001 (Student’s *t*-test). (C) Co-immunoprecipitation of CENP-B with RNF4 or RNF4_DN_ in the presence of the SUMO protease inhibitor N-ethylmaleimide (NEM). (D) Co-immunoprecipitation of CENP-B with RNF4 or RNF4_DN_ in the absence (−) or presence (+) of NEM. CENP-B, RNF4, SUMO-1, SUMO-2, and ubiquitin were detected. (E) Co-immunoprecipitation of RNF4 or RNF4_DN_ with CENP-B in the absence (−) or presence (+) of NEM. CENP-B, RNF4, SUMO-1, and SUMO-2 were detected. Arrows indicate remaining CENP-B signal after stripping of the membranes. (F) Up: WB of CENP-B ectopically co-expressed with RNF4wt or RNF4_DN_. Middle: Quantification of CENP-B (graph and values) and actin (values) amounts in each lane (relative amounts). Down: Quantification of CENP-B mRNA

If RNF4 is involved in the turnover of CENP-B at centromeres then it should incrementally take part in the depletion of CENP-B from the centromeres whenever the CENP-B protein amount is artificially reduced for example by siRNA (Fig. 6A for the synopsis of the experiment). Transfection of a CENP-B-specific siRNA significantly decreased the amount of CENP-B at centromeres in contrast to RNF4-specific siRNA (Fig. 6Bii and iii, and C lanes 3 and 4). Co-transfection of CENP-B and RNF4 siRNAs restored the signal and amount of CENP-B at centromeres (Fig. 6Biv and C lane 5). This was not indirectly due to a differential depletion of the CENP-B mRNA because RT-qPCR did not detect any significant difference in CENP-B mRNA amount when CENP-B siRNA was used alone or together with the RNF4 siRNA (Fig. 6D, data normalized on the actin housekeeping gene). If CENP-B stability is dependent on a SUMOylation/ubiquitination pathway, then E3 SUMO ligases should be involved in that process. Protein inhibitor of activated STATs (PIASs) proteins are known for their SUMOylation activity (Johnson and Gupta, 2001) and have been connected to the centromere activity in yeasts and/or mammalian cells (Azuma et al., 2005; Dasso, 2008; Rytinki et al., 2009; Xhemalce et al., 2004). We then performed similar experiment than above but this time by co-depleting PIAS1, 2, and 4 proteins together with CENP-B (Fig. 7A for the synopsis of the experiment). Similarly to the above results, the inactivation of the PIASs together with CENP-B restored the CENP-B signal at centromeres (Fig. 7Biv and C compare lanes 4 and 5). Altogether, these data demonstrated that RNF4 acted as a SUMO-targeted ubiquitin ligase (STUbL) controlling CENP-B stability through a SUMOylation/ubiquitination and proteasome degradation process involving PIAS proteins.

**Figure 6.**
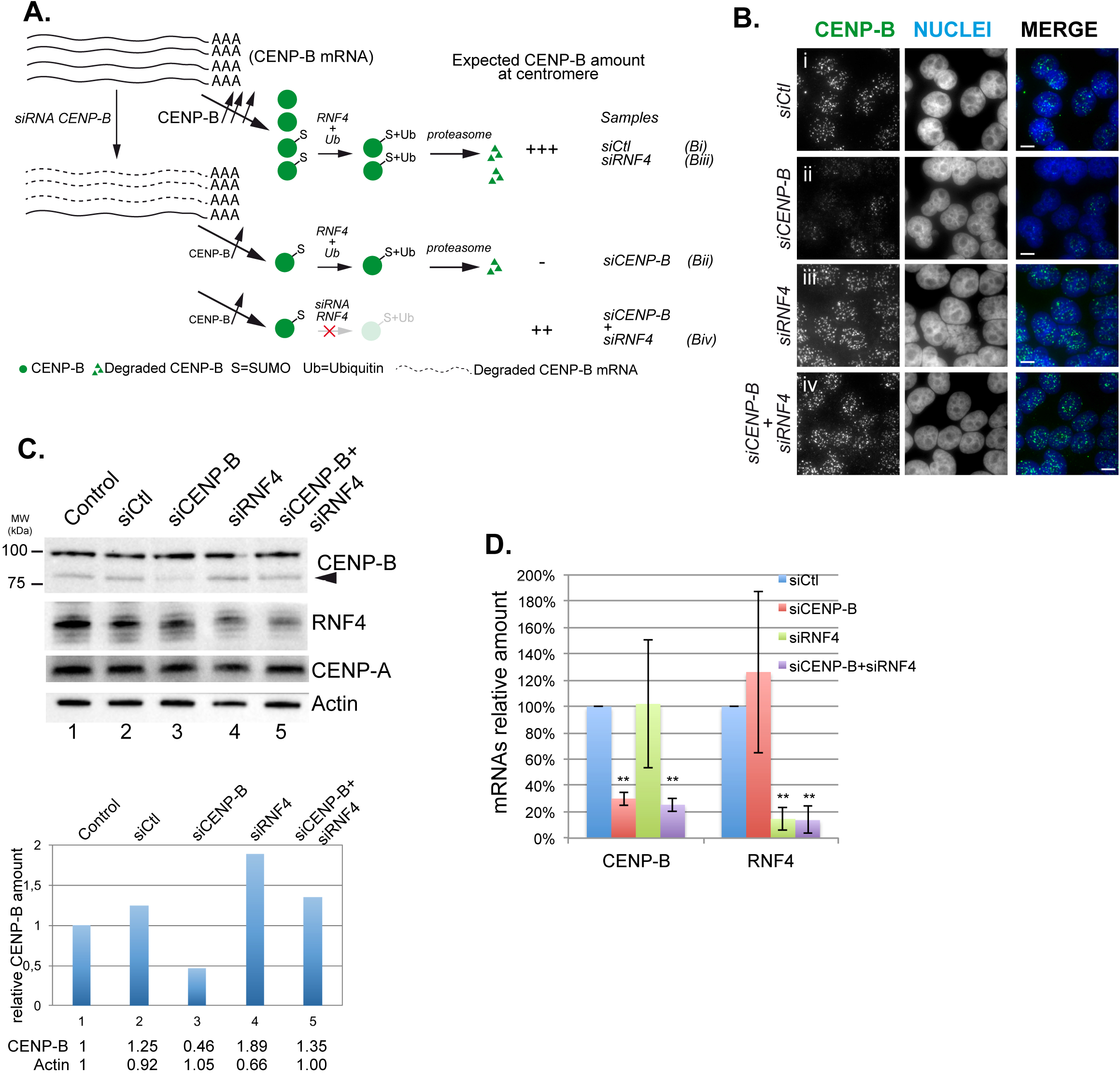
RNF4 is involved in the control of CENP-B stability at centromeres. (A) Synopsis of the experiment. (B) Immunofluorescence to detect endogenous CENP-B in HeLa cells treated with a control siRNA (siCtl) or siRNAs targeting CENP-B, RNF4, or both. (C) Up: WB of endogenous CENP-B, RNF4, or CENP-A of HeLa cell extracts not transfected (Control) or transfected with control siRNA (siCtl), siRNAs targeting CENP-B, RNF4, or both. Actin was used as a loading control. Down: Quantification of CENP-B (graph and values) and actin (values) amounts in each lane (relative amounts). (D) Real Time-quantitative PCR to detect CENP-B or RNF4 mRNA in HeLa cells transfected with control siRNA (siCtl) or siRNAs targeting CENP-B, RNF4, or both. Data were normalized on the actin housekeeping gene. The results show means of three independent experiments (± standard deviation; SD). ** p ≤ 0.01 (Student’s *t*-test).

## Discussion

Centromeres are major chromosome domains essential for the development and survival of all eukaryotes, from yeast to mammals. Yet, their structure and regulation remain largely unknown, in part, due to their complexity. Centromeric DNA is composed of repetitive satellite DNA arrays (called alphoid DNA in humans), themselves forming higher order repeats, which in humans varies from one chromosome to another, making them difficult to experimentally analyze. At the nucleoprotein level, centromeres are packaged into chromatin incorporating a specialized histone variant (CENP-A). In addition, the presence of a set of histone modifications defines the so-called “centrochromatin.” During interphase, tens of proteins interconnect to form the platform that enables the formation of the kinetochore structure to which microtubules attach during mitosis.

CENP-B is one of the first proteins described as a major constituent of the centromere present during the entire cell cycle (Earnshaw and Rothfield, 1985; Earnshaw et al., 1987a; Earnshaw et al., 1987b). CENP-B is a conserved protein in all eukaryotes, which suggests a major role in intra-and inter-species evolution. Despite this, it is understudied partly because previous studies showed that it was not necessary for mouse development, at least until birth (Hudson et al., 1998; Kapoor et al., 1998; Perez-Castro et al., 1998). CENP-B is required for de novo assembly of centromeres with CENP-A-containing chromatin, which confers to CENP-B, at least in some specific contexts, an essential role in the formation of functional centromeres (Okada et al., 2007). In its absence, a higher rate of chromosome mis-segregation is observed in both human and mouse cells, and in fibroblasts from CENP-B null mice, which suggests a conserved role for CENP-B in maintaining chromosomal instability at low levels, preventing aneuploidy and tumorigenesis (Fachinetti et al., 2015, Morozov et al., 2017). A recent study demonstrated the requirement of CENP-B for faithful chromosome segregation following CENP-A depletion from kinetochores, suggesting a major role for CENP-B in maintaining centromere strength during mitosis (Hoffmann et al., 2016). In addition, uncontrolled CENP-B binding to ectopically integrated alphoid DNA promotes both the addition of the H3K9me3 heterochromatin marker and DNA methylation, which were shown to antagonize the formation of CENP-A chromatin (Nakashima et al., 2005; Okada et al., 2007; Okamoto et al., 2007).

In the present study, we demonstrated that CENP-B can be SUMOylated, and that SUMOylation plays a major role in its dynamics at centromeres. Our *in vitro* data show K402 as a major contributor to the SUMOylation of CENP-B. K402 is not required for the targeting of CENP-B to centromeres because its mutation does not affect the accumulation of CENP-B at centromeres. Intriguingly, in the context of the inability of CENP-B protein to bind the CENP-B box due to the deletion of its DBD, and thus to accumulate at centromeres (Pluta et al., 1992), K402 mutation and/or inhibition of proteasome activity partially restored the capacity of CENP-BΔDBD to accumulate at centromeres. These data suggested a role for K402 in the combined SUMO/ubiquitin/proteasome-dependent turnover of CENP-B at centromeres. Our FRAP data indeed showed that K402 mutation affected the mobility of centromere-associated CENP-B, unlike nucleoplasmic CENP-B, and was therefore implicated in CENP-B dynamics at centromeres.

A previous study showed that during interphase, CENPs could be exchanged at the centromeres, revealing unexpectedly complex and dynamic changes within the centromere throughout cell cycle progression (Hemmerich et al., 2008). Two different CENP-B populations were described with drastically different centromere residence times based on the G1/S or G2 cell cycle stages. During the former, CENP-B is very dynamic with a rapid exchange rate, whereas in G2 CENP-B is stably bound to the centromere. If CENP-B K402-dependent centromeric turnover is linked to the process of SUMOylation/ubiquitination and proteasome degradation, then both SUMOs and ubiquitin should be detected at centromeres in some specific experimental conditions. This was indeed the case for SUMOs in a subset of cells in interphase when MG132 treatment was applied for a short period of time. This was much more obvious when MG132 was applied to cells ectopically expressing CENP-Bwt, CENP-BΔDBD, or CENP-BΔDBD_K402G, bypassing the steady-state regulation of centromeric CENP-B. The fact that CENP-BΔDBD_K402G still induced the accumulation of SUMOs and ubiquitin at centromeres, although at least four of its SUMOylable lysines were missing, suggested that (i) it could be due to the modification of endogenous CENP-B; (ii) other lysines in CENP-B may be the sites of SUMOylation (see Supplementary Data in Hendriks et al., 2015) on CENP-B K70 SUMOylation); (iii) the accumulation of CENP-B at centromeres induces the SUMOylation and ubiquitination of other centromeric proteins. The use of CENP-B_4K and CENP-BΔDBD_K402G in our *in vitro* experiments showed that although the SUMOylation of CENP-B was greatly diminished in both proteins, some SUMOylation remained, suggesting the possibility of other SUMOylable lysines, at least *in vitro*. The expression of CENP-BΔDBD and CENP-BΔDBD_K402G, unlike CENP-Bwt in MEF CENP-B^−/−^ cells, did not induce the accumulation of SUMOs or ubiquitin at centromeres in cells treated with MG132. These results demonstrate that there exists a direct correlation between the capacity of CENP-B to accumulate at centromeres and the accumulation of SUMOs and ubiquitin, supporting the likelihood of a direct modification of CENP-B by both SUMOs and ubiquitin. This finding highlighted an unanticipated additional level in the complexity of the regulation of interphase centromere dynamics through the control of CENP-B stability by SUMOylation/ubiquitination and proteasomal degradation. That the CENP-B stability could be controlled by a process of SUMOylation/ubiquitination is not unexpected as we previously described that a viral SUMO-targeted ubiquitin ligase (STUbL), namely ICP0 of herpes simplex virus 1, is capable of inducing the proteasomal degradation of several centromeric proteins including CENP-B (Everett et al., 1999; Lomonte et al., 2000; Lomonte and Morency, 2007; Gross et al., 2012). In addition, a recent study suggested that the integrity of centromeres chromatin could be maintained by the deposition of histones through a CENP-B SUMO-dependent interaction with the histone H3.3 chaperone DAXX (Morozov et al., 2017).

We found that RNF4 and the E3 SUMO ligases, PIASs, cooperate to maintain the CENP-B level at centromeres. RNF4 alone, when over-expressed, was able to deplete CENP-B from centromeres, which suggests a role for this STUbL as an E3 ubiquitin ligase involved in the proteasomal degradation of CENP-B. Given the propensity of CENP-B to be SUMOylated *in vitro* and *in vivo*, and the CENP-B-dependent accumulation of SUMOs at centromeres in proteasome inhibitor-treated cells, we inferred that E3 SUMO ligases could also be involved. SUMO in general and PIASs in particular play critical roles during mitosis (Dasso, 2008). We show that simultaneous inactivation of PIAS1, 2(x), and 4(y) significantly reduced siRNA-induced CENP-B depletion. The requirement for the simultaneous inactivation of 3 PIASs to observe an effect on the reversion of CENP-B depletion is most likely due to redundant PIAS activities. Indeed, PIAS4(y)^−/−^ mice are viable, fertile, and morphologically normal (Roth et al., 2004), suggesting that other PIASs may compensate for the lack of PIAS4(y). Moreover, mutations in yeast genes encoding SUMO ligases do not lead to strong mitotic phenotypes, suggesting that E3 enzymes function redundantly for key substrates (Dasso, 2008). Still, the accumulation of SUMOs at centromeres during the cell cycle is hardly detectable in mammalian cells. This could be potentially explained by the presence of SUMO proteases that, in a very dynamic process, could deSUMOylate substrates to maintain them in a non-SUMOylated form compatible with their stability and activity. In mammalian cells, SUMOylated CENP-I and HP1α are substrates for the SUMO proteases SENP6 and −7, respectively (Mukhopadhyay et al.; 2010 Maison et al., 2012).

All these data are summarized in a working model outlined in Figure 7D. Briefly, following its synthesis, CENP-B is targeted to centromeres where it accumulates. Excess centromeric CENP-B would then be controlled by a SUMOylation/ubiquitination and proteasomal degradation pathway; CENP-B K402 is the keystone of this process. K402 SUMOylation could be induced by PIASs, further activating RNF4. RNF4 then induces the ubiquitination of SUMOylated CENP-B, which signals for its proteasomal degradation. SUMO proteases could then be involved in deSUMOylation of CENP-B K402 to maintain the balance of CENP-B at centromeres.

**Figure 7.**
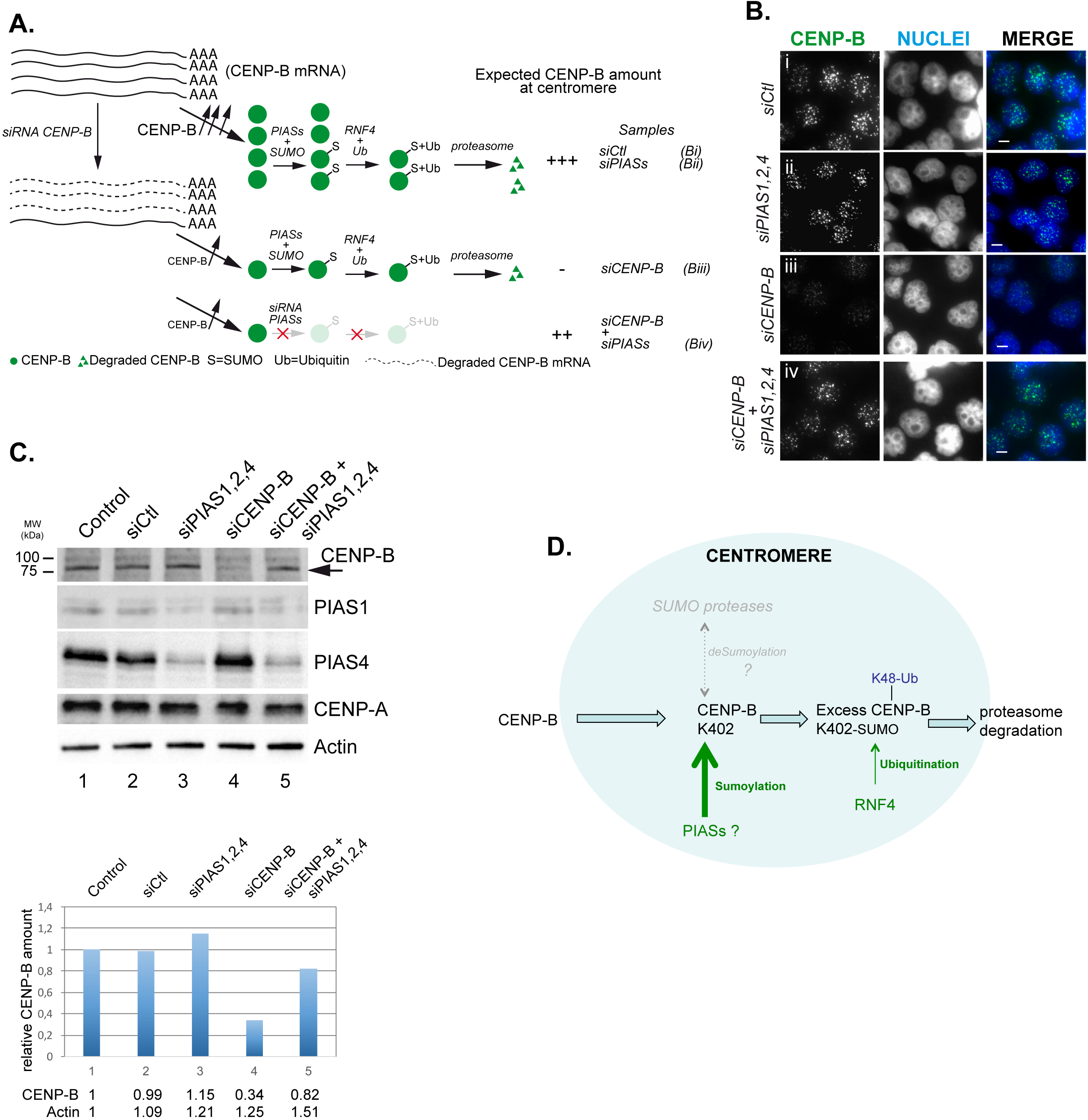
PIASs are involved in the control of CENP-B stability at centromeres. (A) Synopsis of the experiment. (B) Immunofluorescence of endogenous CENP-B in HeLa cells transfected with control siRNA (siCtl), siRNA targeting CENP-B, pooled siRNAs targeting PIAS1, 2, or 4, or CENP-B and PIAS1, 2, or 4. (C) Up: WB of endogenous CENP-B, PIAS1, PIAS4, or CENP-A of HeLa cell extracts not transfected (Control) or transfected with control siRNA (siCtl), siRNA targeting CENP-B, pooled siRNAs targeting PIAS1, 2, or 4, or CENP-B and PIAS1, 2, or 4. Actin was used as a loading control. Down: Quantification of CENP-B (graph and values) and actin (values) amounts in each lane (relative amounts). (D) Model of CENP-B SUMOylation and ubiquitination-dependent turnover at centromeres. Explanations are provided in the Discussion. In grey are hypothetical contributors/activities whose implications have not been demonstrated in the present study.

Both recent and older studies revealed essential CENP-B structural and functional roles at centromeres in the initial formation of a functional locus, as well as their long-term activity in the maintenance of genome stability (Ohzeki et al., 2002; Okada et al., 2007; Fachinetti et al., 2015; Hoffmann et al., 2016). CENP-B acts at the epigenetic level by inducing heterochromatin markers, such as trimethylation of H3K9 (H3K9me3) and CpG methylation (Okada et al., 2007). A mix of euchromatin-and heterochromatin-associated histone markers were described as part of centromere chromatin (Sullivan and Karpen, 2004; Schueler and Sullivan, 2006; Nakano et al., 2008; Mravinac et al., 2009; Stimpson and Sullivan, 2011; Ohzeki et al., 2012; Bergmann et al., 2012; Erliandri et al., 2014). It was suggested that the balance between these markers must be tightly controlled to maintain a chromatin landscape compatible with centromere activity (Nakano et al., 2008). A dual role for CENP-B in centromeric satellite DNA chromatin marker acquisition has been suggested, with CENP-B involved in both CENP-A and H3K9me3 chromatin assembly (Okada et al., 2007; Fachinetti et al., 2015; Morozov et al., 2017). It was shown that a reduction in CENP-A chromatin assembly directly correlates with high levels of H3K9me3 and CpG hypermethylation in alphoid DNA (Okada et al., 2007). Moreover, mouse and human centromeric satellite DNA possess distinctive clusters of CENP-A and H3K9me3 chromatin (Guenatri et al., 2004; Lam et al., 2006; Martens et al., 2005; Nakashima et al., 2005). These features led to the suggestion that CENP-B could be involved in the clustering of centromeric chromatin in major interspersed domains containing either CENP-A nucleosomes and no CpG methylation of centromeric satellite DNA or H3K9me3 nucleosomes and CpG methylation (Okada et al., 2007). An additional impact of CENP-B on the regulation of the timing of replication of alphoid centromere sequences in association with specific histone modification markers has been proposed (Erliandri et al., 2014). The imbalance of the amount CENP-B in one region or another in the alphoid sequence may induce improper chromatin marker acquisition leading to the deregulation of centromere activity.

Among the three CpG potentially methylated sites found in alphoid DNA, two are located in the CENP-B box, and methylation of these sites prevents the binding of CENP-B, which suggests that CENP-B does not occupy all the available CENP-B boxes present in centromeric satellite arrays (Tanaka et al., 2005a; Okada et al., 2007). Additionally, Nap1, a histone chaperone involved in nucleosome assembly by promoting proper DNA binding of core and linker histones, inhibits the non-specific binding of CENP-B to alphoid DNA (Tachiwana et al., 2013). Indeed, CENP-B DBD is highly basic, which confers on CENP-B the propensity to non-specifically bind to the negatively charged DNA phosphate backbone, similar to histones. It thus makes sense that cellular mechanisms are required to “clean” centromeric chromatin from an excess of non-specifically bound CENP-B. Our study brings to light an additional mechanism involved in controlling the amount of CENP-B at the centromeres to maintain centromere function over cell generations. Any failure in that process could lead to centromere inactivation, mitotic defects, and associated genetic instabilities.

## Materials and Methods

### Cells and plasmids

HeLa (ATCC^®^ CCL2™), MEF CENP-B^+/+^ and CENP-B^−/−^ cells were grown at 37°C in Glasgow Modified Eagle (GMEM) or DMEM medium complemented with 10% fetal bovine serum, L-Glutamin (1% v/v), 10 units/ml of penicillin, 100 mg/ml of streptomycin. For MG132 treatments cells were incubated in medium containing 2.5 μM MG132 for five hours.

pcDNA-HA-CENP-B, pcDNA-HA-RNF4, p3X-FLAG-CENP-B, and p3X-FLAG-RNF4 plasmids were constructed by cloning in a pcDNA3.1-HA (LOMONTE et al., 2004) or p3X-FLAG (Sigma) vector, the CENP-B or the RNF4 cDNAs retrieved from a Hela cDNA library.

CENP-B mutants, and RNF4_DN_ (C132/135S)-expressing plasmids were obtained by site-directed mutagenesis using the QuickChange^®^ kit (Agilent). CENP-B K28, K58, K76 and K402 were mutated in P28, G58, G76 and G402 respectively. Multiple lysine-mutated CENP-B sequences were obtained by sequential site-directed mutagenesis.

PET28b-6xHis-CENP-Bwt plasmid was first constructed by insertion of the CENP-Bwt ORF. PET28b-6xHis-CENP-B_Kmut plasmids were then obtained either by sub-cloning from the pcDNA-CENP-B_Kmut plasmids or by site-directed mutagenesis. PET28b-6xHis-FLAG-CENP-Bwt and pET28b-6xHis-FLAG-CENP-B_Kmut plasmids were constructed by in frame insertion of a 69nt fragment containing the 3x-Flag and isolated from the p3xFLAG-CMV10 plasmid, between the 6xHis and the ATG of the CENP-Bwt and mutant ORFs or by sub-cloning. PET28b-6xHis-FLAG-PML was obtained by sub-cloning of PML cDNA in the pET28b-6xHis-FLAG vector.

pLVX-6xHis-SUMO-1 vector was kindly provided by Dr Ben Hale (Institute of Medical Virology, University of Zürich).

pEGFPC1-CENP-Bwt, pEGFPC1-CENP-B_K402G, pEGFPC1- CENP-BΔDBD, pEGFPC1-CENP-BΔDBD_K402G plasmids were obtained by a double digestion of the corresponding pcDNA-HA-CENP-B plasmids and subcloning of the fragments.

All plasmids were purified and verified by sequencing.

### HIS-tagged protein pull-down assays

HeLa cells (2.4 × 10^6^ cells/dish) were transfected with the appropriate plasmids. Twenty four hours post-transfection, cells were washed once with cold phosphate saline buffer (PBS) containing 20mM N-Methylmaleimide (NEM) before adding 4 mLs of denaturing sample buffer (6M GuHCL, 100mM Na2HPO4, 100mM NaH2PO4, 10mM Tris/HCl pH 8.0, 10mM imidazole, ß-mercaptoethanol 5mM, benzonase 250U/mL). After 20min lysis on ice, the solution was sonicated using a probe-sonicator. The debris were pelleted and the supernatant was transferred in a new tube before adding 60 μl of Ni-NTA agarose resin (Quiagen) pre-equilibrated with denaturing sample buffer, and incubated by rotation for at least 16 hours at 4°C. Beads were spun out of suspension and the resin was transferred into a new tube. The resin was washed once with denaturing sample buffer, then twice with denaturing pH 8.0 wash buffer (8M urea, 100mM Na2HPO4, 100 mM NaH2PO4, 10mM Tris/HCl pH 8.0, 10mM imidazole, ß-mercaptoethanol 5mM), then twice with denaturing pH 6.3 wash buffer (8M urea, 100mM Na2HPO4, 100 mM NaH2PO4, 10mM Tris/HCl pH 6.3, 10mM imidazole, ßmercaptoethanol 5mM). Beads were resuspended in 60 μl of laemmli buffer before boiling for 2 min.

### Immunoprecipitation

HeLa cells were seeded at 1.2 × 10^6^ cells per 100 mm Petri dish. The following day, cells were transfected (Effectene Transfection Reagent; Qiagen) with the adequate plasmids. The following day cells were washed once with cold phosphate saline buffer (PBS) containing 20mM NMethylmaleimide (NEM), scraped off the flask then centrifuged at 1000 X g for 5min. The cell pellet was resuspended in 200μl of a lysis buffer (15 mM Tris-HCl (pH 7.5), 2 mM EDTA, 0.25 mM EGTA, 15 mM NaCl, 0.3 mM Sucrose, 0.5 % Triton X 100 (TX100), 300mM KCl, 20mM NEM). Samples were incubated on ice for 30 min, and centrifuged at 6000 X g for 5min at 4°C to remove all debris. Protease inhibitors phenylmethylsulfonyl fluoride (PMSF) and protease inhibitor cocktail tablets (Roche) were added to the lysis buffers. Cell extracts were incubated with the appropriate antibodies O/N at 4°C. Then 50 μl of dynabeads protein G (Thermoscientific) pre-equilibrated in lysis buffer, were added to the samples and incubation was carried on for one hour. Samples were then centrifuged briefly and washed 5 times with lysis buffer before resuspending the beads in Laemmli buffer then boiled.

### Western blotting and proteins quantifications

Samples from immunoprecipitation and siRNA assays were treated for western blotting according to the protocol described in (Lomonte et al., 2000). Proteins were loaded on a SDS 4-15% polyacrylamide gradient gel, before running, transfer and detection. Quantification of proteins was performed using the Image Lab software associated with the ChemiDoc Imaging system (Biorad).

### SiRNA experiments

Cells were seeded at very low density before siRNAs transfections. Two rounds of siRNAs transfections (Effectene Transfection Reagent; Qiagen) were performed at 48 h intervals with the amounts of siRNAs adjusted to the number of cells following the manufacturer’s protocol. Cells were then treated depending on the subsequent experiment to be performed (WB or IF). The siRNA sequences used were: Control siRNA (siCtl) 5’-UACAGCUCUCUCUCGACCC-3’ (Eurogentec); siRNF4: 5’-GAAUGGACGUCUCAUCGUU-3’ (Galanty et al., 2012); PIAS1 siRNA (siPIAS1): 5’-GGAUCAUUCUAGAGCUUUA-3’ (Galanty et al., 2012); PIAS2 siRNA (siPIAS2): 5’-AAGAUACUAAGCCCACAUUUG-3’ {Yang:2005gf}; PIAS4 siRNA (siPIAS4): 5’-GGAGUAAGAGUGGACUGAA-3’ (Galanty et al., 2012); UBC9 siRNA (siUbc9): 5’-GAAGUUUGCGCCCUCAUAA-3’ (Dharmacon, on target J-004910–06) (Pourcet et al., 2010; Rojas-Fernandez et al., 2014). All these siRNAs have been used in previous studies and analyzed for absence of off-target effects.

### Real time quantitative PCR (RT-qPCR)

Detections and quantifications of FLAG-CENP-B, endogenous CENP-B, endogenous RNF4, and actin transcripts were performed using the QuantiTect SYBR Green RT-PCR kit (Qiagen) with the following primers: endoCENP-B fwd: GGGAGGCCATGGCTTACTTT; endoCENP-B rev: 5’-TTCCAAGTGGAGGATGTGGC-3’; RNF4 fwd: 5’-ACTCGTGGAAACTGCTGGAG-3’; RNF4 rev: 5’-TCATCGTCACTGCTCACCAC-3’; FLAG fwd: 5’-CTACAAAGACCATGACGGTGA-3’; ectoCENP-B rev: 5’-GGATGATCCGTGACTTCTCCC-3’; actin fwd: 5’-CGGGAAATCGTGCGTGACATTAAG-3’; actin rev: 5’-GAACCGCTCATTGCCAATGGTGAT-3’. Data were normalized on the actin housekeeping gene.

### FRAP experiments

Hela cells were plated in 27 mm diameter glass-bottom plastic petri dishes (Dominique Dutscher) at a concentration of 5.10^5^ cells/plate. Cells were transfected with plasmids coding for GFP, GFP-CENP-Bwt, GFP-CENP-B_K402G, GFP-CENP-BΔDBD, or GFP-CENP-BΔDBD_K402G using effectene reagent (Qiagen). Bleaching was performed with the 458, 488 and 514 nm lines of an argon laser (set to 100%) and the 561 nm line of a DPS 561 laser. Five pre-bleach images were collected, followed with a 145 ms bleach pulse on a 3 μm diameter spot. Images were collected every 393ms during 120 sec after simultaneous bleaching of two centromeres per nucleus. Images were acquired with the 488 nm line of the argon laser (set to 1%). Loss of fluorescence due to acquisition was calculated using total fluorescence of the cell, and pre-bleach intensity as reference. Photobleaching did not exceed 5%. Fluorescence intensity of centromeres was background substracted, and corrected for photobleaching due to image acquisition. Fluorescence of each photo-bleached centromere was normalized to the fluorescence before bleaching. FRAP experiments were performed on a scanning confocal microscope LSM 780 (Carl Zeiss, Inc.).

### Fluorescent in situ hybridization (FISH)

The protocol used was derived from Solovei et al. (Solovei et al., 2002). Cells were seeded at 7.5×10^4^ cells per well in Millicell EZ SLIDE 8-well glass (Merck Millipore). Cells were fixed with PFA 2% for 10min., washed twice with PBS, permeabilized for 5 min. with Triton X-100 0.5% in PBS, and then washed twice with 2 X SSC. DNA deproteination was performed by incubation of cells in HCl 0.1M for 5 min. Cells were washed twice with 2X SSC, then incubated 2×10min in a solution 50% formamide/SSC 2X. Ten μl of a hybridization solution (10% dextran, 1X denhardt, 2X SSC, 50% formamide) containing 20 ng of probe (StarFISH human chromosome pan-centromeric paints biotin, Cambio, UK), were put in contact with the cells. Coverslips were sealed with “Rubber cement” until dried out. DNA denaturation of cells and probe was performed for 5min at 80°C, and hybridization was carried out overnight at 37°C.

Cells were washed 3×10min in 2X SSC and 3×10min in 0.2X SSC at 37°C then pre-incubated with 4XSSC/5% milk for 30 min. Streptavidin labeled with Alexa-Fluor 488 diluted in 4X SSC/5% milk was added for 1 H RT. Cells were washed twice with 4X SSC, treated with TX-100 0.1%/2X SSC for 10min, then washed twice with 4X SSC, before staining the nuclei with Hoechst 33238 (Invitrogen). Slides were mounted using Vectashield mounting medium (Vector Laboratories) and stored at +4°C until observation.

### Immunofluorescence and immuno-FISH

For immunofluorescence (IF), cells were seeded at 7.5×10^4^ cells per well in Millicell^®^ EZ SLIDE 8-well glass (Merck Millipore). Cells were treated for IF as described in {Morency:2007bt}. For immuno-DNA FISH, cells were fixed with PFA 2% for 10min, washed twice with PBS, permeabilized for 5 min. with Triton X-100 0.5% in PBS then washed with PBS. Cells were incubated for 1 h with the primary antibody diluted in PBS containing 3% FBS. After 3 washes, the secondary antibody was applied for 1 h at 1/200 dilution. Secondary antibodies were AlexaFluor conjugated (Invitrogen). Following immuno-staining, cells were post-fixed in PFA 1% for 5 min, and DNA-FISH was carried out as described above from the HCl step. Samples were examined either by confocal microscopy (Zeiss LSM 510) or using an inverted CellObserver (Zeiss), and a CoolSnap HQ2 camera from Molecular Dynamics (Ropper Scientific). The data were collected using an alpha Plan-APOCHROMAT 100x/1.46 oil lens. Datasets were processed using LSM 510 software, and then by ImageJ software (Rasband, W.S., ImageJ, U.S., National Institutes of Health, Bethesda, Maryland, USA. http://imageJ.nih.gov/ij/) and Adobe Photoshop.

### Antibodies

The following antibodies were used for this study: *mouse monoclonal* anti-6xHis (Clontech, 631212), anti-human CENP-A (Abcam, ab13939), anti-SUMO-1 (MBL, clone 5B12), anti-SUMO2/3 (MBL, clone 1E7), anti-ubiquitin (Affinity, clone FK2), anti-CENP-B (generous gift from Hiroshi Masumoto (Japan), clone 5E6C1); *rat monoclonal* anti-HA (Roche, clone 3F10); *rabbit monoclonal* anti-SUMO-1 (Abcam, clone Y299), anti-SUMO-2/3 (Cell Signaling, clone 18H8), anti-PIAS1 (Abcam, ab109388); *rabbit polyclonal* anti-Ubc9 (Abcam, ab33044); anti-human CENP-A (Abcam, 33565), anti mouse CENP-A (Upstate, 07-574), anti-GFP (LifeTech), anti-HA (Sigma, H6908), anti-SUMO-1 (Cell Signaling, 4971), anti-SUMO2/3 (Abcam, ab3742), anti-RNF4 (Sigma, 1125), anti-actin (Sigma, A2066), anti-PIAS4 (Abcam, ab58416).

### Statistical analyses

Student’s *t*-tests were performed using Microsoft Excel version 12.2.4 for MAC OS X. The results were calculated with the paired test. Results were considered as significant (*) for a p value ≤ 0,05.

### Supplemental material

Supplemental material contains four figures.

## Acknowledgments

We thank H. Masumoto (Kazusa DNA Research Institute, Chiba, Japan) for providing the MEF CENP-B^−/−^ and ^+/+^ cells and the mAb anti-CENP-B (clone 5E6C1) and the Centre Technologique des Microstructures (CTμ) of the Université Claude Bernard Lyon 1 for the confocal microscopy. PL is a CNRS Research Director.

## Authors contribution

Conceptualization: JEM, PL

Funding acquisition: PL

Investigation: JEM, PT, IE, CC, AC, FC, CB

Methodology: JEM, FC, CB, PL

Supervision: PL

Validation: JEM, FC, CB, PL

Writing – original draft: PL

Writing – review & editing: JEM, AC, FC, CB, PL

## Competing interests

No competing interests declared

## Funding

This work was funded by grants from CNRS (http://www.cnrs.fr), INSERM (https://www.inserm.fr), University Claude Bernard Lyon 1 (https://www.univ-lyon1.fr), French National Agency for Research-ANR (PL, ML, VIRUCEPTION, ANR-13-BSV3-0001-01, http://www.agence-nationale-recherche.fr), LabEX DEVweCAN (PL, CC, ANR-10-LABX-61, http://www.agence-nationalerecherche.fr), La Ligue régionale contre le Cancer and the FINOVI foundation (grant #142690).

## Figure legends

**Figure S1.**
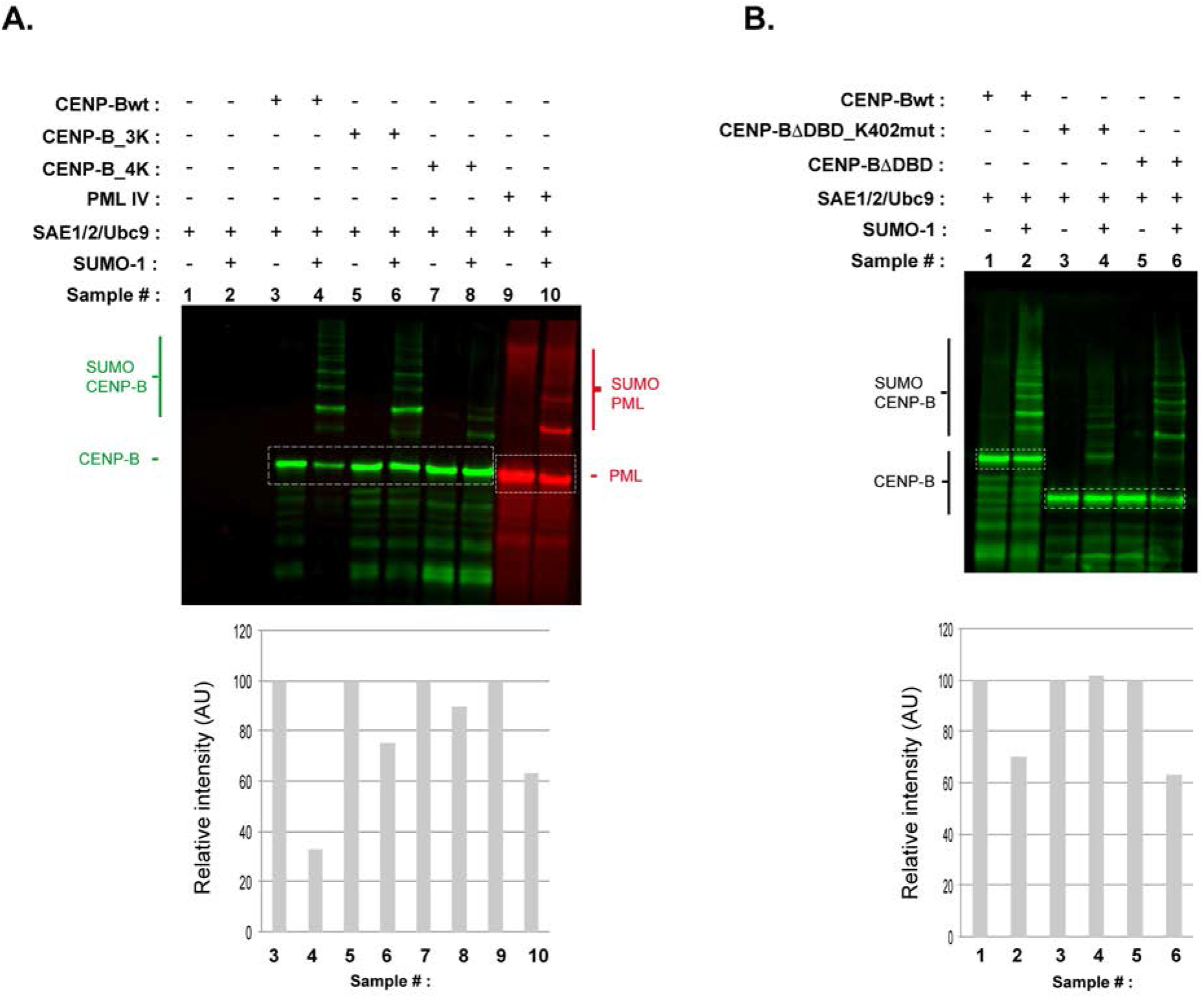
In vitro SUMOylation assays of: (A) 6xHis-FLAG-CENP-B, 6xHis-FLAG-CENP-B_3K, 6xHis-FLAG-CENP-B_4K, and 6xHis-FLAG-PML isoform IV (used as positive control); (B) 6xHis-FLAG-CENP-B, 6xHis-FLAG-CENP-BΔDBD_K402G, and 6xHis-FLAG-CENP-BΔDBD Tops: WB was performed using an anti FLAG antibody for the detection of tagged proteins. LI-COR Infrared Fluorescent Imaging System was used for the detection and quantification of the signals. Bottoms: (A) quantification of the CENP-B wt and mutants, and PML unmodified bands (boxed in the WB image). Unmodified CENP-Bwt signal decreased oppositely to the increase of CENP-Bwt-associated SUMO forms (compare signal relative intensity in boxed area of lanes 3 and 4). CENP-B_3K also showed a decrease in its relative intensity following SUMOylation compared to that of the unmodified protein (compare lanes 5 and 6), suggesting that although these K residues within the DBD do not contribute to the majority of CENP-B SUMOylation they were likely modified; (B) quantification of the CENP-Bwt and truncated CENP-B unmodified bands (boxed in the WB image). Unmodified CENP-B signals decreased oppositely to the increase of CENP-B-associated SUMO forms.

**Figure S2.**
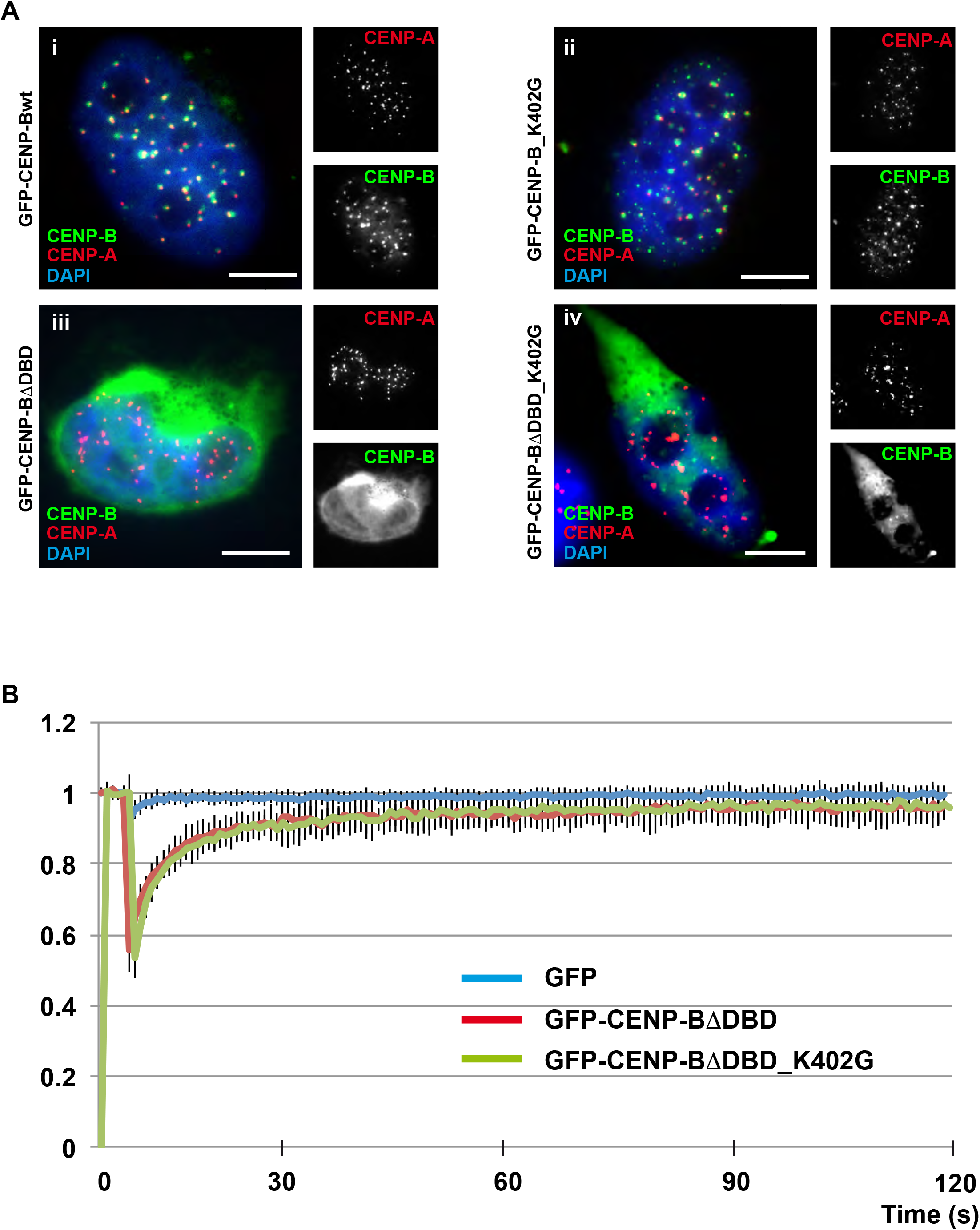
(A) Immunofluorescence to detect GFP-CENP-B (i), GFP-CENP-B_K402G (ii), GFP-CENP-BΔDBD (iii), and GFP-CENP-BΔDBD_K402G (iv) at centromeres in HeLa cells. Centromeres were visualized by the detection of CENP-A. Bars = 5 μm. (B) Fluorescence Recovery After Photobleaching (FRAP) analysis of GFP, GFP-CENP-BΔDBD and GFP-CENP-BΔDBD_K402G mobility in the nucleoplasm. Bleaching was performed in the nucleoplasm using a single bleach spot. GFP was used as a reference of freely diffusing protein.

**Figure S3.**
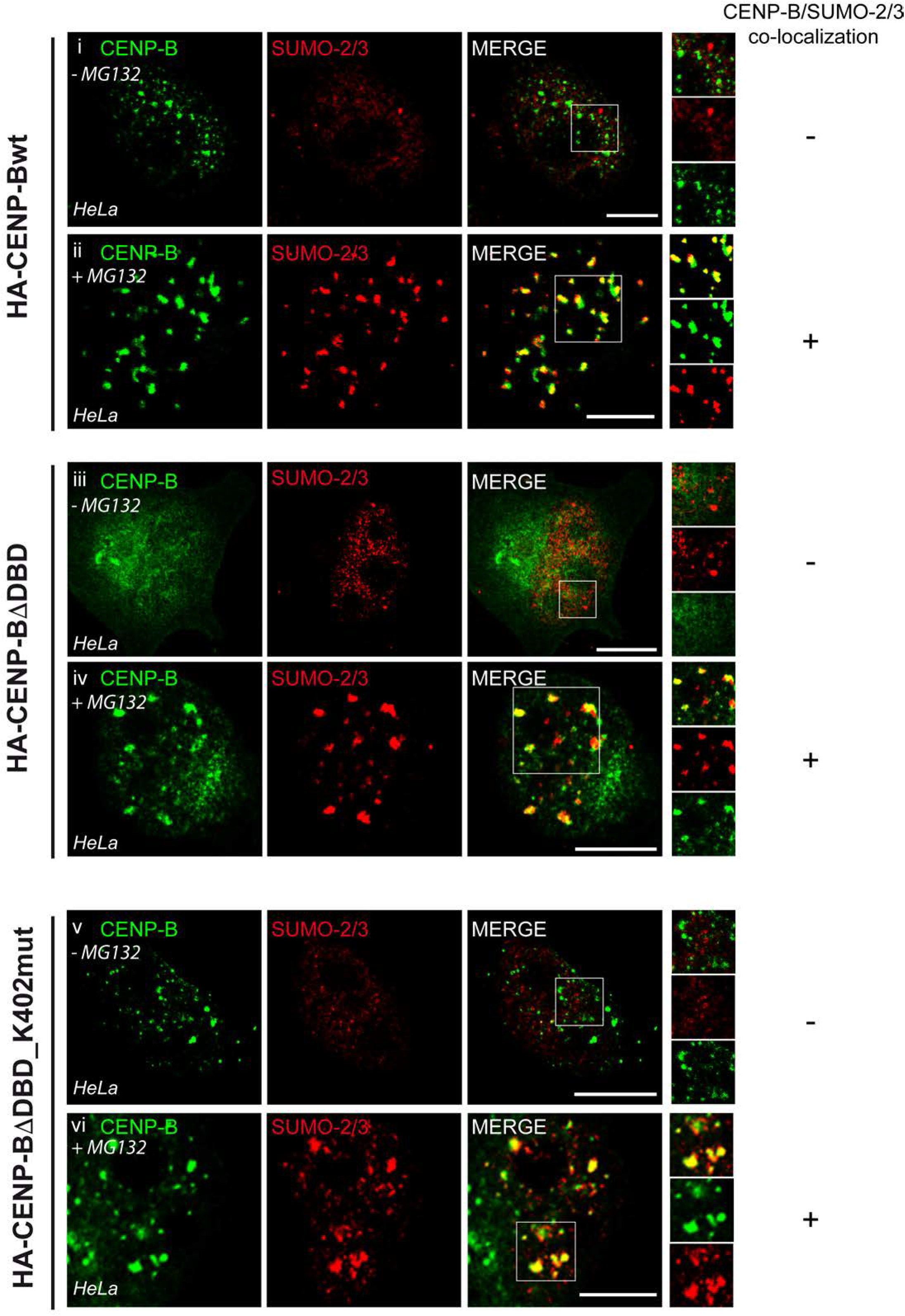
Immunofluorescence to detect SUMO-2/3 at centromeres in HeLa cells expressing HA-CENP-Bwt, HA-CENP-BΔDBD, HA-CENP-BΔDBD_K402G in absence or presence of the proteasome inhibitor MG132. Bars = 5μm. +: indicates that co-localization between CENP-B proteins and SUMO-2/3 could be observed; -: indicates that no co-localization between CENP-B proteins and SUMO-2/3 could be observed.

**Figure S4.**
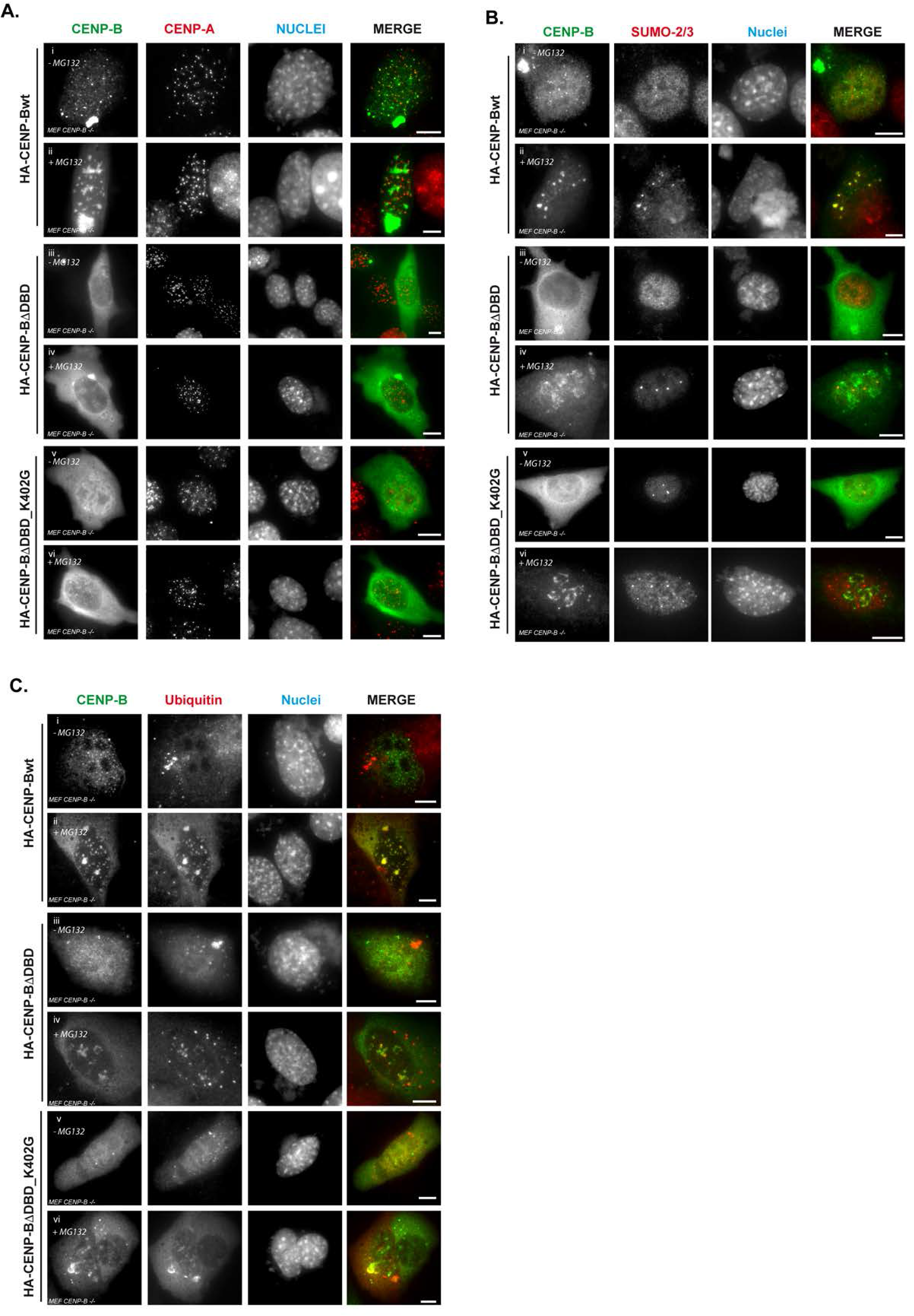
Immunofluorescence to detect CENP-B and CENP-A (A) or SUMO-2/3 (B) or ubiquitin (C) in MEF CENP-B^−/−^ cells expressing HA-CENP-Bwt, HA-CENP-BΔDBD, HA-CENP-BΔDBD_K402G in absence or presence of proteasome inhibitor MG132. Bars = 5μm.

**Table 1:** Centromeric localization of ectopically expressed CENP-B in MEF CENP-B^−/−^ cells

| HA-CENP-B | Description | HA-CENP-B positive cells showing CENP-B co-localization with CENP-A (%) |
| --- | --- | --- |
| _Wt | Wild type CENP-B | 87,9 ± 10.5 |
| _ΔDBD | CENP-B deleted of its 129 first aa containing the DNA binding domain | 0 |
| _K402G | CENP-B mutated on K402 | 90.3 ± 8.1 <sup>(1)</sup> |
| _ΔDBD_K402G | CENP-B deleted of its 129 first aa containing the DNA binding domain and additionally mutated on the K402 | 4.1 ± 2.6 |

<sup>(1)</sup> Although with a similar pattern than CENP-Bwt, the CENP-B\_K402G shows more intense centromeric spots

